# Heparin inhibits cellular invasion by SARS-CoV-2: structural dependence of the interaction of the surface protein (spike) S1 receptor binding domain with heparin

**DOI:** 10.1101/2020.04.28.066761

**Authors:** Courtney J. Mycroft-West, Dunhao Su, Isabel Pagani, Timothy R. Rudd, Stefano Elli, Scott E. Guimond, Gavin Miller, Maria C. Z. Meneghetti, Helena B. Nader, Yong Li, Quentin M. Nunes, Patricia Procter, Nicasio Mancini, Massimo Clementi, Antonella Bisio, Nicholas R. Forsyth, Jeremy E. Turnbull, Marco Guerrini, David G. Fernig, Elisa Vicenzi, Edwin A. Yates, Marcelo A. Lima, Mark A. Skidmore

## Abstract

The dependence of the host on the interaction of hundreds of extracellular proteins with the cell surface glycosaminoglycan heparan sulphate (HS) for the regulation of homeostasis is exploited by many microbial pathogens as a means of adherence and invasion. The closely related polysaccharide heparin, the widely used anticoagulant drug, which is structurally similar to HS and is a common experimental proxy, can be expected to mimic the properties of HS. Heparin prevents infection by a range of viruses if added exogenously, including S-associated coronavirus strain HSR1. Heparin prevents infection by a range of viruses if added exogenously, including S-associated coronavirus strain HSR1. Here, we show that the addition of heparin to Vero cells between 6.25 - 200 μg.ml^−1^, which spans the concentration of heparin in therapeutic use, and inhibits invasion by SARS-CoV-2 at between 44 and 80%. We also demonstrate that heparin binds to the Spike (S1) protein receptor binding domain and induces a conformational change, illustrated by surface plasmon resonance and circular dichroism spectroscopy studies. The structural features of heparin on which this interaction depends were investigated using a library of heparin derivatives and size-defined fragments. Binding is more strongly dependent on the presence of 2-O or 6-O sulphation, and the consequent conformational consequences in the heparin structure, than on N-sulphation. A hexasaccharide is required for conformational changes to be induced in the secondary structure that are comparable to those that arise from heparin binding. Enoxaparin, a low molecular weight clinical anticoagulant, also binds the S1 RBD protein and induces conformational change. These findings have implications for the rapid development of a first-line therapeutic by repurposing heparin as well as for next-generation, tailor-made, GAG-based antiviral agents against SARS-CoV-2 and other members of the *Coronaviridae*.

## Introduction

Heparin, the second most widely used drug by weight globally, is formulated as a polydisperse, heterogeneous natural product. Unfractionated heparin, low molecular weight heparins, such as enoxaparin, and heparinoids are clinically approved as anti-coagulants and anti-thrombotic agents with excellent safety, stability, bioavailability and pharmacokinetic profiles. Crucially, heparin and its derivatives, some of which lack significant anticoagulant activity ^1^, constitute an under-exploited antiviral drug class. This is despite the fact that they exhibit broad-spectrum activity against numerous distinct viruses, including *Coronaviridae* and SARS-associated coronavirus strain HSR1 ^2^, in addition to flaviviruses ^3,4^, herpes ^5^, influenza ^6^ and HIV ^7,8^.

Conventional drug development processes are slow to respond to emerging public health threats, such as the current SARS-CoV-2 coronavirus outbreak, which makes the repurposing of existing drugs a timely and attractive alternative. Heparin, a well-tolerated anticoagulant pharmaceutical, has been used safely in medicine for over 80 years and, alongside its anticoagulant activities, its ability to prevent viral infection, including by members of the *Coronaviridae*, has been described ^2^. The closely related glycosaminoglycan (GAG) member, heparan sulphate (HS), is also known to bind CoV surface proteins and is exploited by coronavirus for its attachment to target cells ^9^. Furthermore, anticoagulant and anti-inflammatory therapy, which are both associated with heparin use, have been shown to be beneficial for COVID-19 management ^10,11^.

Heparan sulphate is a ubiquitous component of the cell surface and the extracellular matrix with a complex non-template driven biosynthesis, involving many enzymes, the consequence of which is a heterogeneous structure. Currently, HS and heparin are known to bind to over 700 proteins ^12^. Heparan sulphate and heparin are linear, sulphated polysaccharides, comprising repeating disaccharide units of a uronic acid and D-glucosamine. They are linked by 1,4 glycosidic bonds and the uronic acid can be either β D-glucuronic acid or its C-5 epimer, α L-iduronic acid (properly, glucuronate and iduronate respectively under physiological conditions). The disaccharide can be subject to several further modifications; iduronate can be O-sulphated at position-2, glucosamine can be N-sulphated, N-acetylated or unmodified, and adorned with O-sulphates at position-6 and, less frequently, at position-3 ^13^.

The anticoagulant properties of heparin arise predominantly from its interaction with serine proteases, particularly antithrombin and the pentasaccharide sequence; GlcNS,6S-GlcA-GlcNS,3S,6S-IdoA2S-GlcNS,6S, has been identified as being most significant in this regard ^14^. On the other hand, its antiviral and anti-inflammatory properties have not been linked to specific sequences enabling strategies to be developed for the production of heparin with tailored activities (i.e., non-anticoagulant but anti-viral) that involve cleaving/disrupting the chain at the antithrombin interacting pentasaccharide, or by globally altering the chemical properties of the chain. Since heparin is a complex mixture, containing polysaccharide chains of varied composition and bearing different biological activities, an alternative approach would be to fractionate the heterogeneous heparin mixture and isolate those fractions with the required activities.

As yet, there are no commercially available medicinal products designed to treat and/or prevent infections associated with the current SARS-CoV-2 coronavirus outbreak. Here, we describe the ability of heparin to inhibit cell invasion and bind the SARS-CoV-2 S1 receptor binding domain (RBD) and investigate the structural requirements and consequences of binding. The results underpin the development of heparin-based therapeutics for Covid-19 and future viral diseases.

## Methods & Materials

### 2.1 Viral invasion assay

Vero cells (ECACC) were plated at 2.5 × 10^5^ cells per well in 24-well plates and incubated with EMEM supplemented with 10% (v/v) foetal calf serum (complete medium). Twenty four hours later, cells were incubated with heparin (from 200 μg.ml^−1^; Celsus Laboratories, Cincinnati, USA) in 300 μL of complete medium 1 hour prior to infection and then incubated with virus solution containing 50 plaque forming units (PFU) of either; Italy/UniSR1/2020 strain (GISAID accession ID: EPI_ISL_413489 ^15^, or, SARS-CoV HSR-1 (EID 2004). After incubation for 1 hour at 37°C, supernatants were discarded and 500 μL of 1% (w/v) methylcellulose (Sigma-Aldrich, Italy) overlay (in complete medium) were added to each well. After 3 days, cells were fixed using a 6% (v/v) formaldehyde: phosphate-buffered saline solution and stained with 1% (w/v) crystal violet (Sigma-Aldrich, Italy) in 70% (v/v) methanol (Sigma-Aldrich, Italy). The plaques were counted under a stereoscopic microscope (SMZ-1500, Nikon).

### 2.2 Recombinant expression of SARS-CoV-2 S1 RBD

Residues 330−583 of the SARS-CoV-2 Spike Protein (GenBank: MN908947) were cloned upstream of a N-terminal 6XHisTag in the pRSETA expression vector and transformed into SHuffle® T7 Express Competent *E. coli* (NEB, UK). Protein expression was carried out in MagicMedia™ *E. coli* Expression Media (Invitrogen, UK) at 30°C for 24 hrs, 250 rpm. The bacterial pellet was suspended in 5 mL lysis buffer (BugBuster Protein Extraction Reagent, Merck Millipore, UK; containing DNAse) and incubated at room temperature for 30 mins. Protein was purified from inclusion bodies using IMAC chromatography under denaturing conditions. On-column protein refolding was performed by applying a gradient with decreasing concentrations of the denaturing agent (6 - 0 M Urea). After extensive washing, protein was eluted using 20 mM NaH_2_PO_4_, pH 8.0, 300 mM NaCl, 500 mM imidazole. Fractions were pooled and buffer-exchanged to phosphate-buffered saline (PBS; 140 mM NaCl, 5 mM NaH_2_PO_4_, 5 mM Na_2_HPO_4_, pH 7.4; Lonza, UK) using a Sephadex G-25 column (GE Healthcare, UK). Recombinant protein was stored at −20°C until required.

### 2.3 Preparation of chemically modified heparin derivatives

All chemically modified heparin polysaccharides (**Table 1**) were synthesised from parental unfractionated porcine mucosal heparin (Mw = 12 kDa; Celsus Laboratories, Cincinnati, USA) as previously described ^6,16^. The veracity of all chemical modifications was ascertained using ^1^H and ^13^C NMR, with chemical shifts compared to TSP (Sigma-Aldrich, UK) as an external reference standard.

**Table 1.** Repeating Structures of heparin derivatives. ^*^Additionally, O-sulphated at positions C-3 of glucosamine and the uronic acid.

|  | R<sub>1</sub> | R<sub>2</sub> | R<sub>3</sub> |
| --- | --- | --- | --- |
| Heparin 1 | -SO<sub>3</sub><sup>-</sup> | -SO<sub>3</sub><sup>-</sup> | -SO<sub>3</sub><sup>-</sup> |
| Heparin 2 | -SO<sub>3</sub><sup>-</sup> | -COCH<sub>3</sub> | -SO<sub>3</sub><sup>-</sup> |
| Heparin 3 | -OH | -SO<sub>3</sub><sup>-</sup> | -SO<sub>3</sub><sup>-</sup> |
| Heparin 4 | -SO<sub>3</sub><sup>-</sup> | -SO<sub>3</sub><sup>-</sup> | -OH |
| Heparin 5 | -OH | -COCH<sub>3</sub> | -SO<sub>3</sub><sup>-</sup> |
| Heparin 6 | -SO<sub>3</sub><sup>-</sup> | -COCH<sub>3</sub> | -OH |
| Heparin 7 | -OH | -SO<sub>3</sub><sup>-</sup> | -OH |
| Heparin 8 | -OH | -COCH<sub>3</sub> | -OH |
| Heparin 9 * | -SO<sub>3</sub><sup>-</sup> | -SO<sub>3</sub><sup>-</sup> | -SO<sub>3</sub><sup>-</sup> |

### 2.4 Secondary structure determination of SARS-CoV-2 S1 RBD by circular dichroism spectroscopy

The circular dichroism (CD) spectrum of the SARS-CoV-2 S1 RBD in PBS was recorded using a J-1500 Jasco CD spectrometer (Jasco, UK), Spectral Manager II software (JASCO, UK) and a 0.2 mm pathlength, quartz cuvette (Hellma, USA) scanning at 100 nm.min^−1^ with 1 nm resolution throughout the range 190 - 260 nm. All spectra obtained were the mean of five independent scans, following instrument calibration with camphorsulfonic acid. SARS-CoV-2 S1 RBD was buffer-exchanged (prior to spectral analysis) using a 5 kDa Vivaspin centrifugal filter (Sartorius, Germany) at 12,000 g, thrice and CD spectra were collected using 21 μl of a 0.6 mg.ml^−1^ solution in PBS, pH 7.4. Spectra of heparin (porcine mucosal heparin), its derivative and oligosaccharides were collected in the same buffer at approximately comparable concentrations, since these are disperse materials. Collected data were analysed with Spectral Manager II software prior to processing with GraphPad Prism 7, using second order polynomial smoothing through 21 neighbours. Secondary structural prediction was calculated using the BeStSel analysis server ^17^.To ensure that the CD spectral change of SARS-CoV-2 S1 RBD in the presence of porcine mucosal heparin did not arise from the addition of the heparin alone, which is known to possess a CD spectrum at high concentrations ^18,19^ a difference spectrum was analysed. The theoretical, CD spectrum that resulted from the arithmetic addition of the CD spectrum of the SARS-CoV-2 S1 RBD and that of the heparin differed from the observed experimental CD spectrum of SARS-CoV-2 S1 RBD mixed with heparin. This demonstrates that the change in the CD spectrum arose from a conformational change following binding to porcine mucosal heparin.

### 2.5 Surface Plasmon Resonance determination of SARS-CoV-2 S1 RBD binding to unfractionated heparin

Human FGF2 was produced as described by Duchesne *et al.* ^20^. Porcine mucosal heparin was biotinylated at the reducing end using hydroxylamine biotin (ThermoFisher, UK) as described by Thakar *et al.* ^21^. Heparin (20 μl of 50 mg mL^−1^) was reacted with 20 μl hydroxylamine-biotin in 40 μl 300 mM aniline (Sigma-Aldrich, UK) and 40 μl 200 mM acetate pH 6 for 48 h at 37 °C. Free biotin was removed by gelfiltration chromatography on Sephadex G25 (GE LifeSciences, UK).

A P4SPR, multi-channel Surface Plasmon Resonance (SPR) instrument (Affinté Instruments; Montréal, Canada) was employed with a gold sensor chip that was plasma cleaned prior to derivatization. A self-assembled monolayer of mPEG thiol and biotin mPEG was formed by incubating the chip in a 1 mM solution of these reagents at a 99:1 molar ratio in ethanol for 24 hrs ^22^. The chip was rinsed with ethanol and placed in the instrument. PBS (1X) was used as the running buffer for the three sensing and a fourth background channel at 500 μl.min^−1^, using an Ismatec pump. Twenty micrograms of streptavidin (Sigma, UK; 1 ml in PBS) were injected over the four sensor channels. Subsequently, biotin-heparin (1 ml) was injected over the three sensing channels.

Binding experiments used PBS with 0.02% Tween 20 (v/v) as the running buffer. The ligand was injected over the three sensing channels, diluted to the concentration indicated in the figure legends at 500 μl.min^−1^. Sensor surfaces with bound FGF2 were regenerated by a wash with 2 M NaCl (Fisher Scientific, UK). However, this was found to be ineffectual for SARS-CoV-2 S1 RBD. Partial regeneration of the surface was achieved with 20 mM HCl (VWR, UK) and only 0.25 % (w/v) SDS (VWR, UK) was found to remove the bound protein. After regeneration with 0.25 % (w/v/) SDS, fluidics and surfaces were washed with 20 mM HCl to ensure all traces of the detergent were removed. Background binding to the underlying streptavidin bound to the mPEG/biotin mPEG self-assembled monolayer was determined by injection over the control channel. Responses are reported as the change in plasmon resonance wavelength, in nm and for the three control channels represent their average response.

### 2.6 Prediction of SARS-CoV-2 S1 heparin binding domain

The methods employed to predict heparin binding sequences within the SARS-CoV-2 S1 RBD are described in Rudd *et al.* ^23^. A brief description of the method follows. In combination, Ori *et al.* ^24^ and Nunes *et al.* ^12^ identified 786 heparin binding proteins. In this study, 776 heparin binding proteins, from the previously identified list, were decomposed into amino acid sequences containing no less than 3 residues with at less one basic amino acid. To simplify the problem, the heparin binding proteins were fragmented into sequences containing the following combination of amino acids; BX, BXA, BXS, BXP, BXAS, BXAP and BXPS (where, B = basic, X = hydrophobic, A = acidic, P = polar and S = special). These sequences were compared using a metric based on the Levenshtein distance, a measure of the similarity between strings of characters, which is defined as the minimum number of insertions, deletions or substitutions that need to be applied to the characters to alter one sequence so that it matches the other. Sequences with a similarity score of greater than 0.7 (70% similarity) were considered highly conserved. These sets of highly conserved basic amino acid containing sequences found in heparin binding proteins were then used to identify possible heparin binding sequences within SARS-CoV-2 S1, utilizing the same Levenshtein similarity cut-off of 0.7.

## Results

### 3.1 SARS-CoV-2 viral plaque forming assays

Progeny infectious virion production was measured in culture supernatants using standard plaque forming assays on Vero cells with and without pre-treatment of incubated heparin (from 200 μg.ml^−1^, one hour prior to infection) with both a historical SARS-CoV strain HSR-1 and the recent SARS-CoV-2 strain, Italy/UniSR1/2020. Significant decreases were observed in the number of PFU upon heparin treatment for both SARS-CoV and SARS-CoV-2, with the later demonstrating significantly higher levels of inhibition (80%) compared to that of SARS-CoV **(Figure 1**).

**Figure 1.**
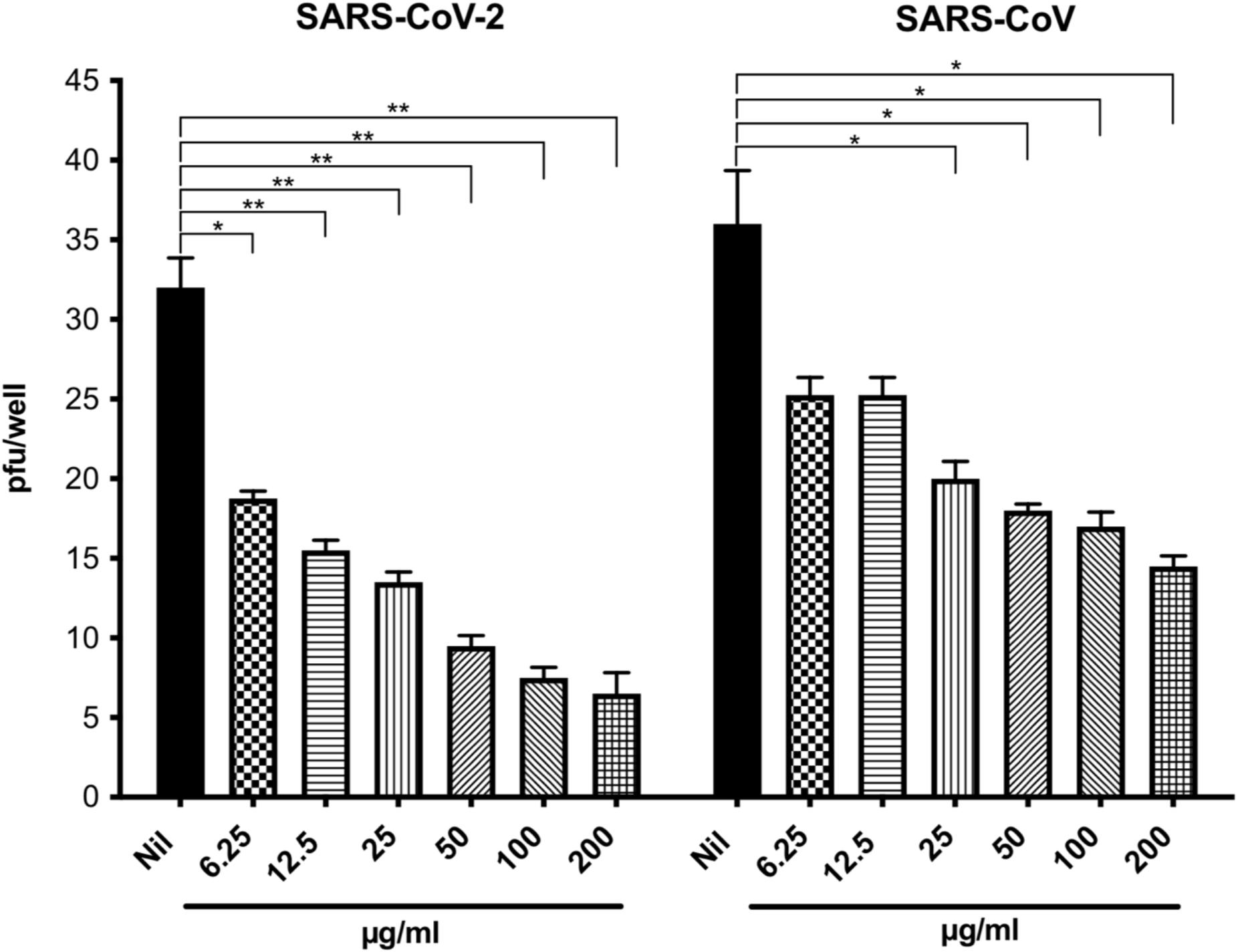
The heparin-mediated inhibition of SARS-CoV-2 viral invasion of Vero cells. The effect of unfractionated porcine mucosal heparin (from 200 μg.ml^−1^) added one hour before the infection of Vero cells with 50 PFU of SARS-CoV-2 or SARS-CoV. Nil represents no treatment. The results are expressed as number of PFU per well and represent the mean ± SD of quadruplicate cultures. The *p* value was calculated using the Mann-Whitney U test.

### 3.2 Surface Plasmon Resonance binding studies

FGF2, a well-characterised heparin-binding protein was used to confirm the successful functionalization of the three sensing channels with biotin-heparin. Injection of 1 ml 100 nM FGF2 over the sensing channels elicited a significant response (**Figure 2A**, injection between the blue and red arrow). However, 100 nM FGF2 elicited no response in the control channel, functionalized solely with streptavidin (**Figure 2A**). The bound FGF2 was removed by a wash with 2 M NaCl, as previously for the IASys optical biosensor ^16^.

**Figure 2.**
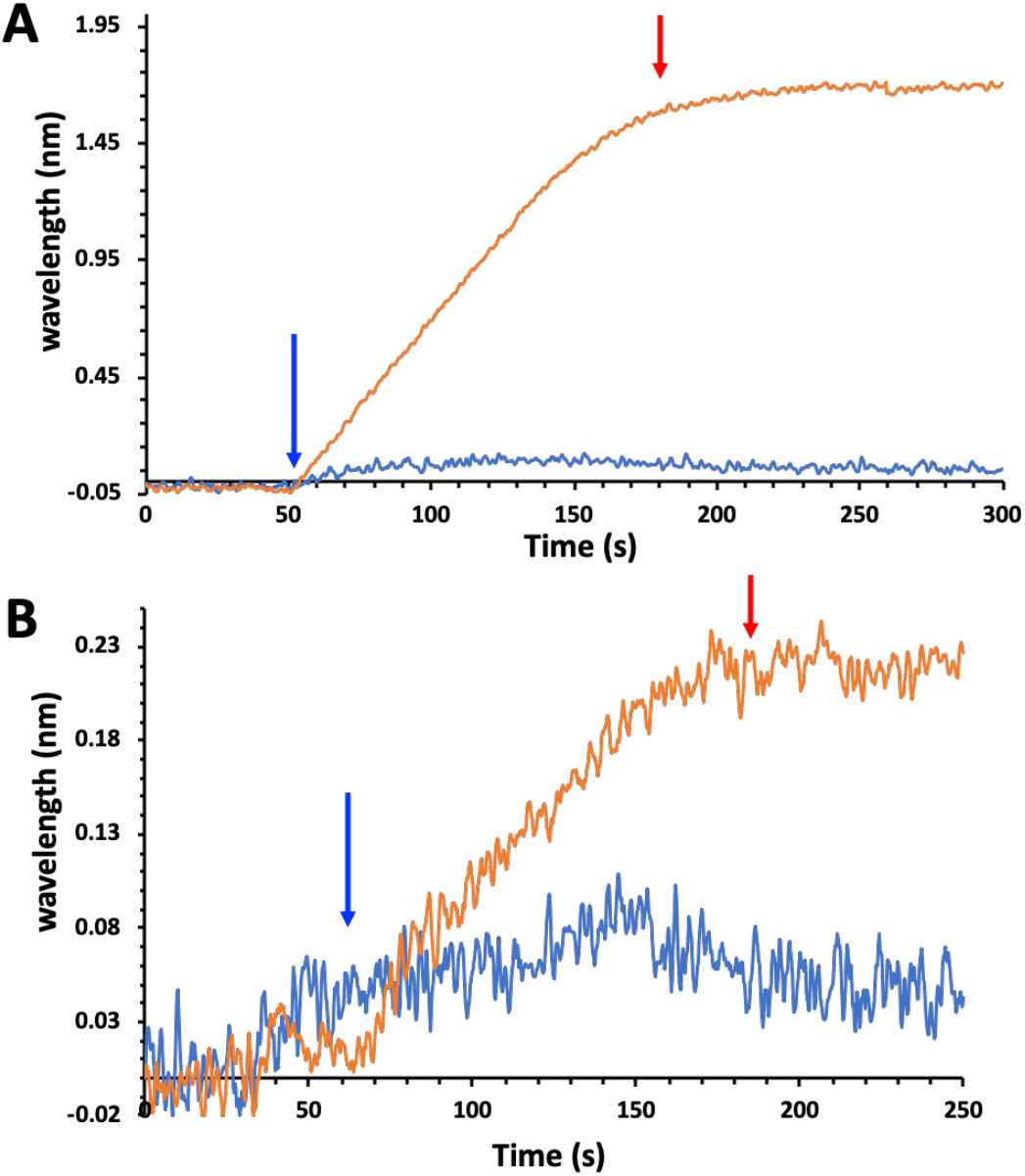
Interaction of FGF2 and 100 nM SARS-CoV-2 S1 RBD with immobilised heparin. Reducing end biotinylated heparin 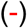 was immobilised on a streptavidin functionalised P4SPR sensor surface (no biotin-heparin 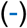 control). PBS running buffer flow rate was 500 μl.min^−1^. The data for the three sensing channels are reported as an average response 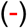. The start of protein injections are indicated by blue arrows and the return of the surface to running buffer (PBST) by red arrows. (**A**) Injection of 100 nM FGF2. (**B**) Injection of 100 nM SARS-CoV-2 S1 RBD protein.

When 100 nM SARS-CoV-2 S1 RBD was injected over the three sensing channels, there was an increase in signal, of binding (**Figure 2B** injection start blue arrow). At the end of the injection the system returned to running buffer (PBST, red arrow). There was no dissociation. This is common in surface measurement, particularly in flow systems, due to the extensive boundary layer of liquid enabling rebinding of the analyte and often causes a substantial underestimation of the dissociation rate, k_off_, which can be remedied by the addition of soluble ligand ^25^. The injection of 100 nM SARS-CoV-2 S1 RBD over the control channel, functionalised with just streptavidin, showed a small increase in response (**Figure 2B**) of the order of 10% of that seen in the measurement channel. Repeated measurements indicate that this is the maximum level of background binding. These data demonstrate that 100 nM SARS-CoV-2 S1 RBD binds specifically to heparin immobilised through its reducing end and does not interact to a major extent with the underlying streptavidin/ethyleneglycol surface. It should be noted that when biotin-heparin is anchored to the streptavidin layer, as in the measurement channels, such background binding will be reduced, since less of the underlying surface is exposed. This is illustrated in the competition experiments, in which soluble heparin was able to completely abrogate the binding of SARS-CoV-2 S1 RBD to the surface (**Figure 3A**).

**Figure 3.**
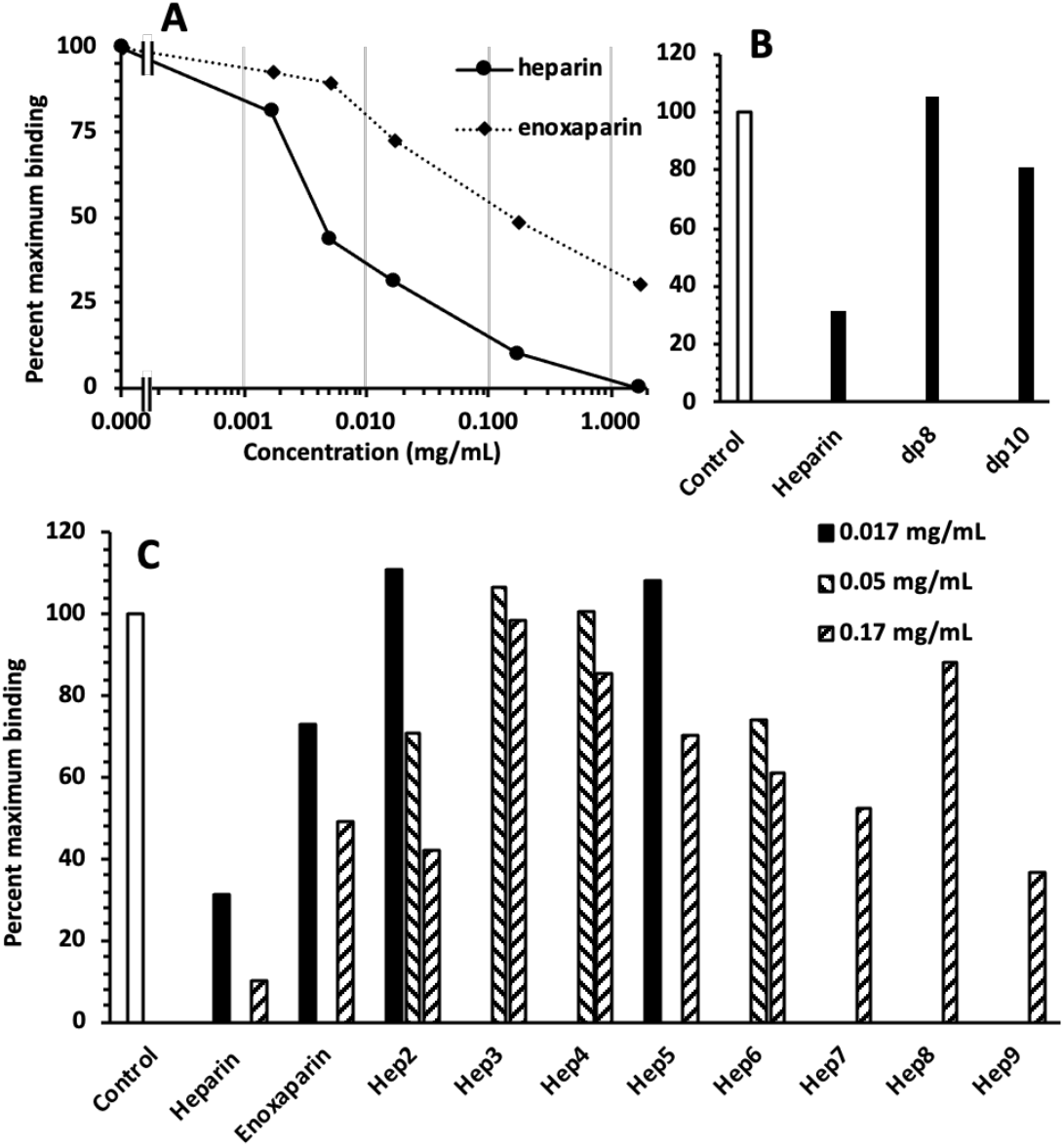
Competition for SARS-CoV-2 S1 RBD binding to immobilised heparin with model heparin-derived oligo- and polysaccharides. 100 nM SARS-CoV-2 S1 RBD was injected onto the surface in the presence or absence of the indicated concentration of heparin-derived oligo- and polysaccharides. Since the SARS-CoV-2 S1 RBD does not dissociate appreciably when the system returns to PBST, this is when the response was measured, to avoid any confounding effects of differences of refractive index between samples. A control measurement (100 nM SARS-CoV-2 S1 RBD alone) was performed before each competition and used to calculate the percentage of maximum binding. This ensured that small changes over time in the responsiveness of the surface did not cofound the analysis. (**A**) Competition for 100 nM SARS-CoV-2 S1 RBD binding to immobilised heparin by heparin and by enoxaparin. (**B**) Competition for 100 nM SARS-CoV-2 S1 RBD binding to immobilised heparin by 0.17 mg.ml^−1^ heparin-derived dp 8 and dp 10, with the corresponding value for heparin from panel (A) shown to aid comparison. (**C**) Competition for 100 nM SARS-CoV-2 S1 RBD binding to immobilised heparin by a panel of orthogonally chemically desulphated heparins (**Table 1**).

In a first set of competition experiments, immobilised 100 nM SARS-CoV-2 S1 RBD was exposed to heparin or enoxaparin at the indicated concentration. The response when the surface returned to running buffer, PBST, was then measured. This is appropriate, since there is no appreciable dissociation of the heparin-bound SARS-CoV-2 S1 RBD when the surface returns to PBST (**Figure 2B**) and it ensures that any differences in refractive index of samples do not interfere with the measurement. At 0.0017 mg.ml^−1^ heparin, a small reduction in the binding of SARS-CoV-2 S1 RBD was observed (**Figure 3A**) and, as the concentration of heparin increased, the binding of SARS-CoV-2 S1 RBD decreased in a dose-dependent manner. SARS-CoV-2 S1 RBD binding was completely abrogated by 1.7 mg.ml^−1^ heparin. Enoxaparin was also found to inhibit the binding of SARS-CoV-2 S1 RBD, but on a weight basis was less potent than heparin (**Figure 3A**). Thus, a small inhibition of binding was observed at 0.017 mg/mL Enoxaparin and the maximal inhibition observed with this polysaccharide was 70% at 1.7 mg.ml^−1^. These data indicate that heparin is ~30-fold more potent an inhibitor of the interaction of SARS-CoV-2 S1 RBD with immobilised heparin than Enoxaparin. We then examined whether short oligosaccharides could inhibit SARS-CoV-2 S1 RBD binding. A heparin-derived octasaccharide (dp 8) was without effect, but a decasaccharide (dp10) at 0.17 mg.ml^−1^ showed a modest inhibition of SARS-CoV-2 S1 RBD binding to immobilised heparin. Whether these data reflect a true requirement for a longer structure for SARS-CoV-2 S1 RBD binding or instead, reflect a selective reduction in the presence of particular binding structures in the polysaccharide as a consequence of the processes (β-elimination for enoxaparin and partial enzymatic heparin degradation for oligosaccharides), remains to be determined.

We next tested the ability of a library of heparins that had been selectively modified (**Table 1**) to inhibit SARS-CoV-2 S1 RBD binding. These showed different levels of inhibitory activity, depending on their pattern of sulphation. Of the singly desulphated heparins, heparin 2, which is *N*-desulphated/*N*-re-acetylated, showed inhibitory activity at 0.05 mg.ml^−1^ and 0.17 mg.ml^−1^, whereas heparin 3 and heparin 4, which are 2-*O* and 6-*O* desulphated, respectively, had no detectable inhibitory activity (**Figure 3C**). This suggests that SARS-CoV-2 S1 RBD has a preference for tracts of saccharides that are 2-*O* and 6-*O* sulphated. Interestingly, the doubly desulphated heparins (heparin 5, heparin 6, heparin 7) possessed inhibitory activity, albeit lower than the native heparin (**Figure 3C**). Since the sulphation of the polysaccharide has a marked effect on its conformation ^16,18,19,26^ these data suggest that the SARS-CoV-2 S1 RBD may have a preference for a particular spatial arrangement of charged groups. Completely desulphated heparin (heparin 8) has no inhibitory activity, indicating that ionic interactions with sulphate groups make an important contribution to the interaction of the SARS-CoV-2 S1 RBD with the polysaccharide. Per-sulphated heparin (heparin 9) inhibited most strongly of the heparin derivatives, but not as effectively as native heparin. Again, this indicates the likely importance of the conformation of the polysaccharide for SARS-CoV-2 S1 RBD binding, since sulphation on all available hydroxyls renders the polysaccharide more rigid and restrict is conformational flexibility ^26^.

### 3.3 Secondary structure determination of SARS-CoV-2 S1 RBD protein by circular dichroism spectroscopy

Circular dichroism (CD) spectroscopy in the far UV region (190 - 260 nm) detects conformational changes in protein secondary structure that occur in solution and can infer binding by an added ligand. Such secondary structural changes can be quantified using spectral deconvolution ^27^. SARS-CoV-2 S1 RBD underwent conformational change in the presence of heparin (**Figures 4** and **5**), consisting of an increase in α-helix content of 1.5% and a decrease in global β-sheet of 2.1%. The observed changes demonstrate that the SARS-CoV-2 S1 RBD interacts with heparin in aqueous conditions of physiological relevance. A chemically modified heparin derivative with the predominant repeating disaccharide structure; IdoA-GlNAc,6S, was able to induce closely comparable secondary structural changes in the SARS-CoV-2 S1 RBD as heparin (**Figure 5, A - C**). Analysis by CD spectroscopy of the role of chain length for heparin-derived oligosaccharides (**Figure 6, A - D**) revealed that a hexasaccharide fraction was able to induce similar conformational changes to heparin.

**Figure 4.**
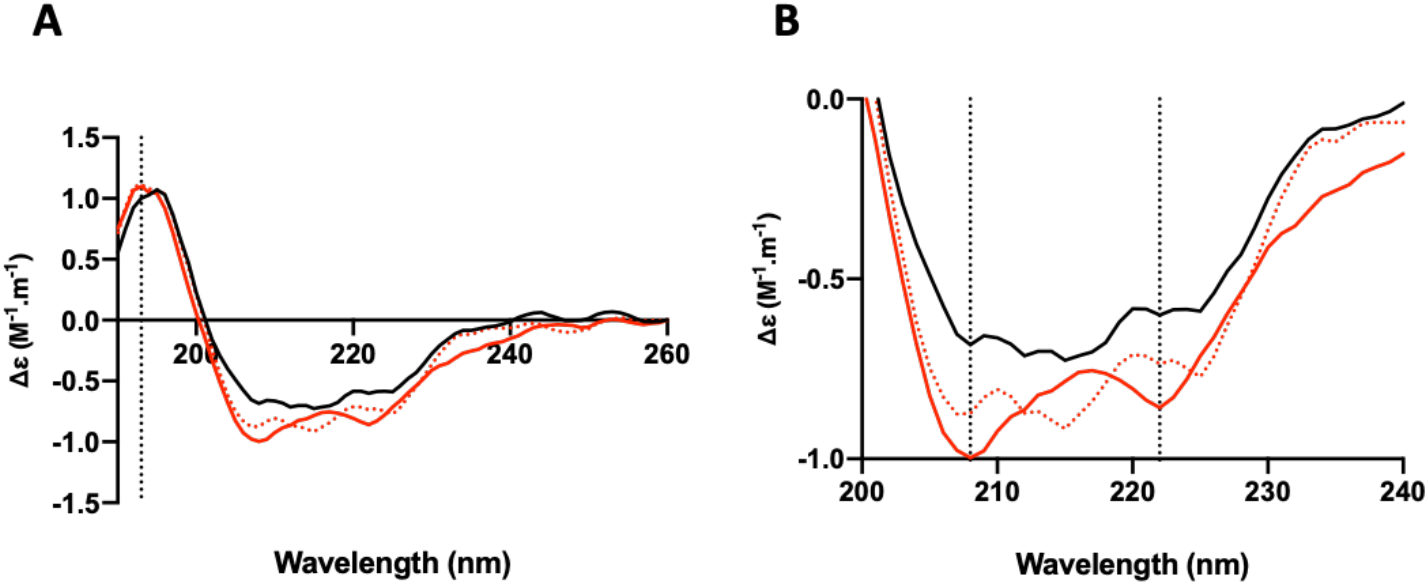
The conformational change of the SARS-CoV-2 S1 RBD observed in the presence of heparin by circular dichroism (CD) spectroscopy. (**A**). Circular dichroism spectra (190 - 260 nm) of nCovS1RBD alone (black solid line) and heparin (red solid line) in PBS, pH 7.4. The red, dotted line represents the sum of the two individual spectra. The dotted vertical line indicates 193 nm. (**B**) Details of the same spectra expanded between 200 and 240 nm. Vertical dotted lines indicate 222 nm and 208 nm.

**Figure 5.**
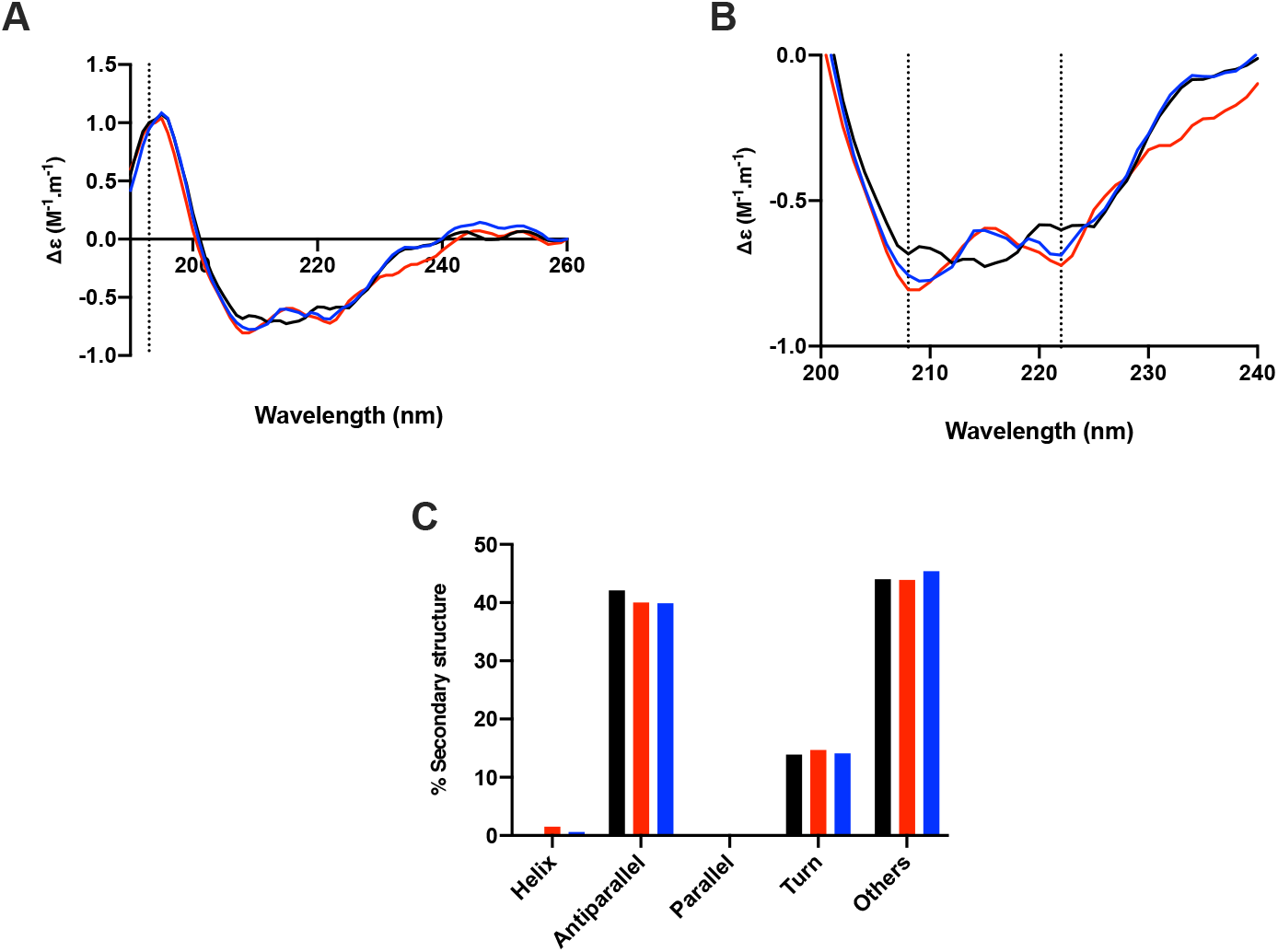
The conformational change of the SARS-CoV-2 S1 RBD observed in the presence of a chemically modified heparin derivative by circular dichroism (CD) spectroscopy. (**A**). Circular dichroism spectra (190 - 260 nm) of nCovS1RBD alone (black) with heparin (red) and a chemically modified derivative, heparin 5, with the predominant repeating disaccharide structure; –IdoA2OH-GlcNAc6S– (blue) in PBS, pH 7.4. The vertical dotted line indicates 193 nm* (**B**). The same spectra expanded between 200 and 240 nm. Vertical dotted lines indicate 222 nm and 208 nm*. (**C**) Secondary structure content analysed using BeStSel for nCovS1RBD. α helical secondary structure is characterized by a positive band at ~193 nm and two negative bands at ~208 and ~222 nm (analysis using BeStSel was performed on smoothed data between 190 and 260 nm.

**Figure 6.**
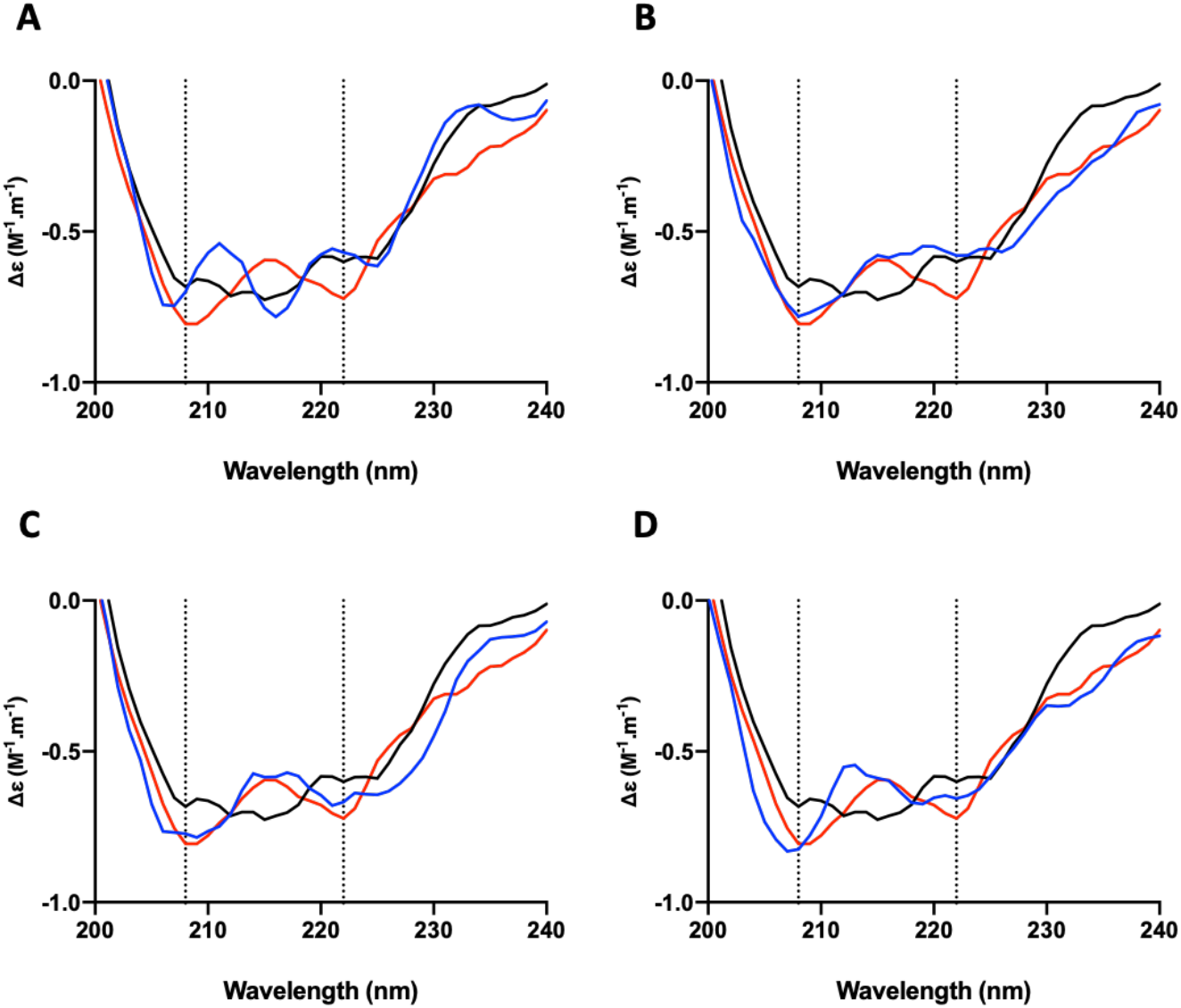
The conformational change of the SARS-CoV-2 S1 RBD observed in the presence of a size-defined heparin oligosaccharides by circular dichroism (CD) spectroscopy. Circular dichroism spectra between 200 and 240 nm of nCovS1RBD in PBS, pH 7.4 alone (black), with heparin (red) and PMH-derived, size defined oligosaccharides (blue): (**A**) tetrasaccharide, (**B**) hexasaccharide, (**C**) octasaccharide and (**D**) decasaccharide. Vertical dotted lines indicate 222 and 208 nm.

### 3.4 Heparin binding site analysis

Basic amino acids are known to dominate the binding between proteins and heparin. With that in mind, primary sequence analysis of the expressed protein domain and analysis of the modelled SARS-CoV-2 S1 RBD structure (**Figure 7**) were conducted and indicated that there are several potential heparin binding sites and, importantly, these patches of basic amino acids are exposed on the protein surface.

**Figure 7.**
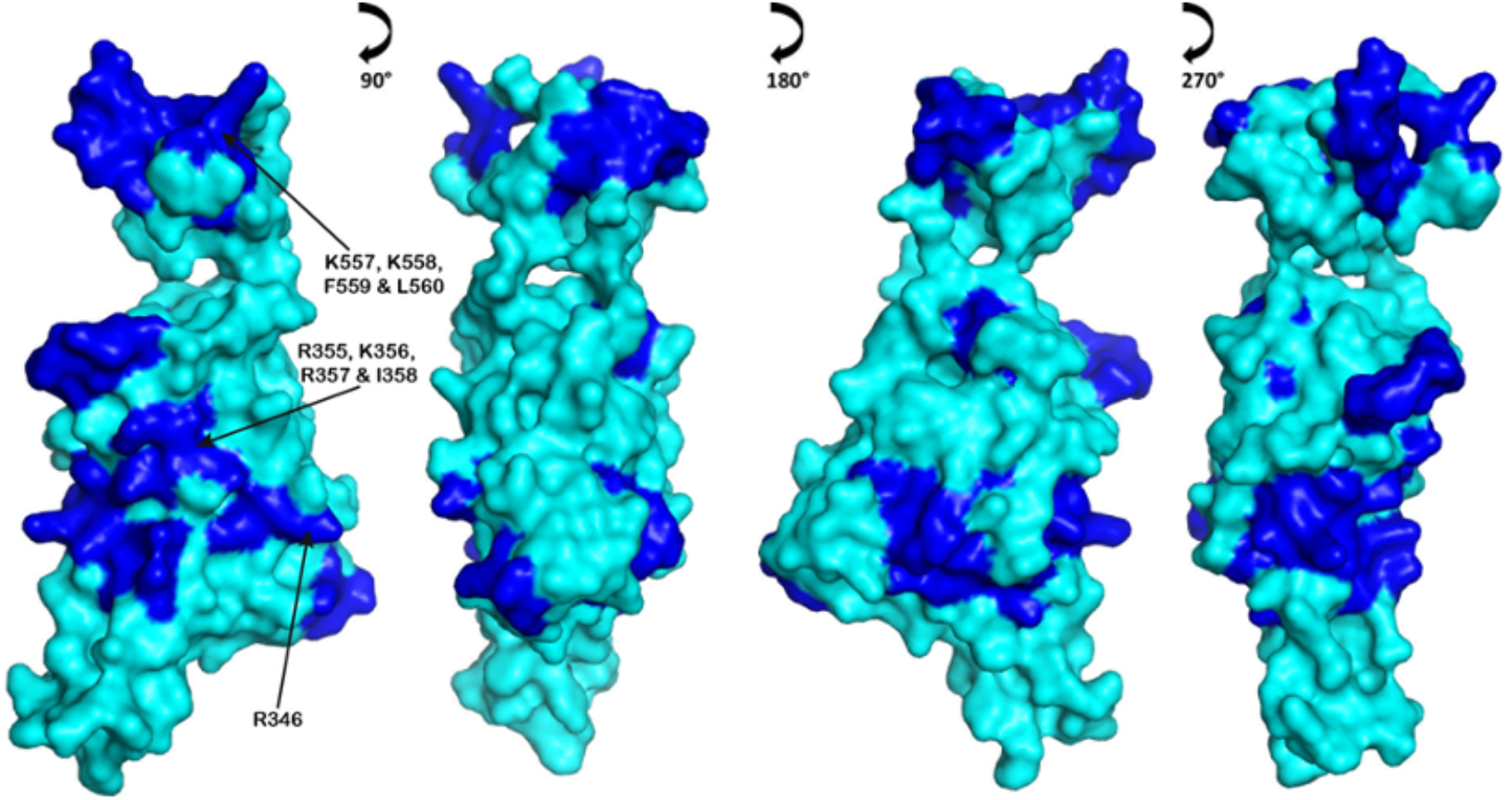
SARS-CoV-2 S1 RBD protein model. Basic amino acids that are solvent accessible on the surface are indicated (dark blue); these form extensive patches. Sequences with the highest normalised count (**Tables 2** and **3**) are highlighted. R346 is also shown as it indicates a potential heparin-binding gain of function mutation (T346R) from the Bat-RaTG13.

Analysis of the RBD sequence for potential heparin binding sites employing a metric based on the Levenshtein distance (a measure of the similarity between two sequences) found that the basic amino acids sequences within SARS-CoV-2 S1 RBD were similar to 278 sequences found in 309 heparin binding proteins. The predicted heparin binding basic amino acid contain sequences are shown in **Tables 2** and **3**. Basic amino acids with a normalised count greater than 0.5 were found at; RKR 355-356 (0.941), LVK 533-535 (1.00), KK 557-558 (0.544) and R 557 (0.688). There were also a number of possible secondary sites of interaction (a normalised count great than 0.3); R 346 (0.368), R 403 (0.386), K 417 (0.309) and H 519 (0.305). One consequence of the ability of SARS-CoV-2 S1 RBD to interact with HS is that it provides a route for adhering to cell surfaces, enabling invasion. Conversely, this region also interacts with the orthodox receptor of the spike protein ACE2 (SARS-CoV-2 S1 RBD 436-529), suggesting that heparin and its derivatives interfere with the interactions between the virus, via these residues. Furthermore, most of the identified sequences, with 3 exceptions (TKLN, GKIADYNYKLP and PYRVVVL), are exposed on the protein surface and available for binding.

**Table 2.**
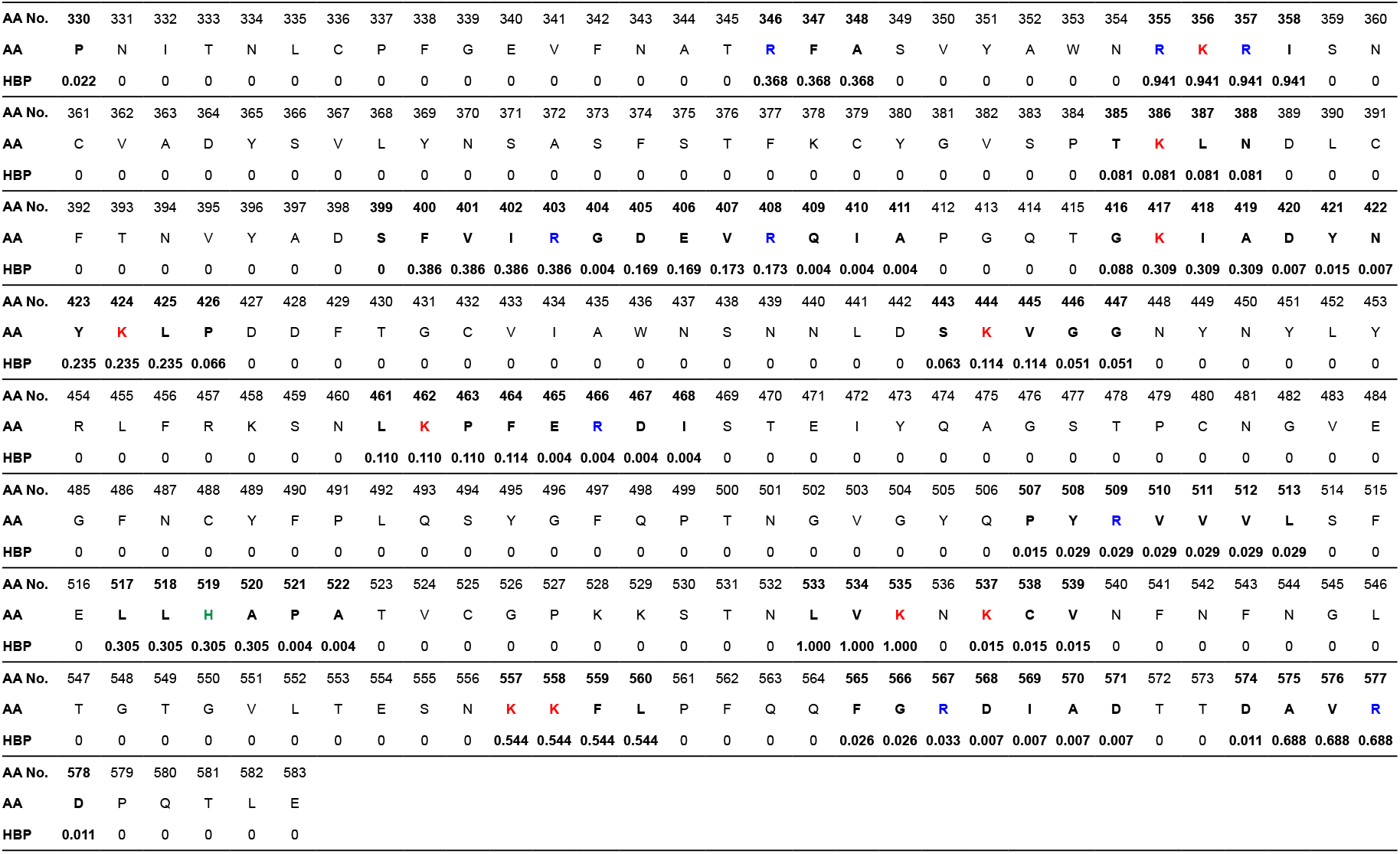
Sequence analysis of SARS-CoV-2 S1 RBD (330-583). The amino acid sequence, number and normalized count for similar sequences in the SARS-CoV-2 RBD as found in a library of 776 heparin binding proteins. The higher the value, the more often the short heparin binding sequence was identified among the set of heparin binding proteins. Basic amino acids within regions of high similarity are identified; arginine (blue), lysine (red) and histidine (green). AA No; amino acid number. AA; amino acid identity. HBP; relative frequency of sequence among heparin binding proteins.

**Table 3.**
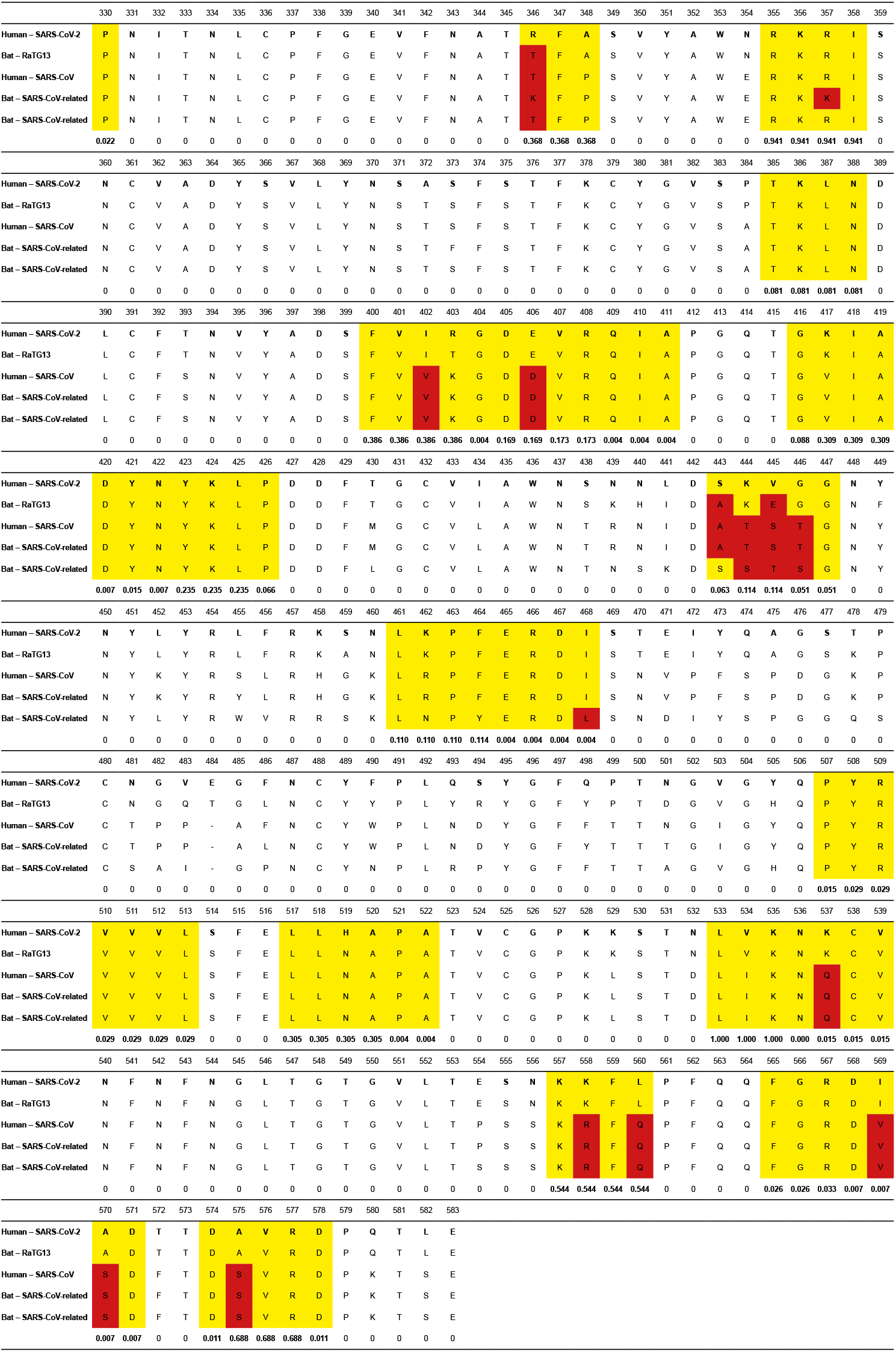
Predicted heparin binding domains in SARS-CoV-2 S1 RBD (330-583). The amino acid sequence, number and normalized count for similar short, heparin binding sequences in the SARS-CoV-2 RBD found in a library of 776 heparin binding proteins. In addition to SARS-CoV-2. The table also contains the aligned sequences from similar viruses, SARS-CoV and related bat viruses. The predicted heparin binding regions are shown in yellow and variations within those regions between SARS-CoV-2 and the related viruses are highlighted in red.

Studying SARS-CoV-2 Spike protein structure and behaviour in solution is an important step in the development of effective therapeutics against SARS-CoV-2. Here, the ability of the SARS-CoV-2 S1 RBD to bind pharmaceutical heparin, studied using spectroscopic techniques, showed that SARS-CoV-2 S1 RBD binds to heparin and that, upon binding, a significant conformational change is induced. Moreover, moieties of basic amino acid residues that are common constituents of heparin binding domains, and accessible to solvent on the SARS-CoV-2 S1 RBD surface, form a continuous patch ((R454, R457, K458, K462, R466, R346, R355, K356 and R346 in **Figure 7**), suitable for heparin binding.

Comparison of the RBD amino acid sequence (residues 330-583) with an extensive library of sequences from known heparin binding proteins (**Tables 2** and **3**) based on the *Levenstein Distance,* which provides a measure of the degree of similarity between the sequences, suggests that wider areas of the RBD surface may be available (**Figure 7**) for binding to host cell surface glycosaminoglycan polysaccharides. These may relate to the propensity for the virus to select particular species, individuals and age groups, since their GAG composition is known to vary. There is evidence of a potential heparin-binding gain of function mutation (T346R) from the Bat-RaTG13 (**Figure 7**).

## Discussion and Conclusion

The rapid spread of SARS-CoV-2 represents a significant challenge to global health authorities and, since there are no currently approved drugs to treat, prevent and/or mitigate its effects, repurposing existing drugs is both a timely and appealing strategy. Heparin, a well-tolerated anticoagulant drug, has been used successfully for over 80 years with limited and manageable side effects. Furthermore, heparin belongs to a unique class of pharmaceuticals for which effective antidotes are available, making it safer to use.

Glycosaminoglycans are present on almost all mammalian cells and this class of carbohydrate is central to the strategy employed by the *Coronaviridae* to attach to host cells. Heparin has previously been shown to inhibit SARS-associated coronavirus strain HSR1 cell invasion ^2^ and its potential against SARS-CoV-2 represented an attractive possibility.

The observation that heparin is able to inhibit invasion by SARS-CoV-2, in a dose-dependent manner at concentrations from 6.25 - 200 μg.ml^−1^, up to 80% in Vero cells is a significant finding that offers one potential route for prophylaxis, as well as high-lighting the capacity of this class of biological macromolecule to intervene in microbial, particularly viral, infection. This ability, in addition to useful properties as anticoagulants and moderators of inflammation, remains under-studied and under-exploited, but offers potential for intervention in many microbial interactions, not least those relating to emerging viral diseases.

The dependence on particular structural features of the heparin scaffold, the resulting changes in conformation of the spike protein RBD and the prediction of heparin-binding features all provide evidence that the interaction with heparin involves a level of structural specificity. They also suggest that heparin, a complex mixture of polysaccharides, contains a range of activities that have the potential of being separated, with or without further modification, to provide diverse derivatives that provide a range of properties that can be tailored to suit a variety of clinical circumstances.

Heparin and its derivatives are amenable to routine parenteral administration through currently established routes and additionally, directly to the respiratory tract via nasal administration, using nebulised heparin, which would be unlikely to gain significant access to the circulation. Thus, the anticoagulant activity of heparin which can, in any event, be engineered out through suitable chemical modification, would not pose a problem. Importantly, such a route of administration would be suitable for prophylaxis and also for patients under mechanical ventilation ^28^.

Together, these data support the use of GAG-derived pharmaceuticals against SARS-associated coronavirus. Furthermore, this study supports the repurposing of heparin and its derivatives as antiviral agents, to provide a rapid countermeasure against the current SARS-CoV-2 outbreak and future emerging viral diseases.

---

**Supplementary Figure S1:**
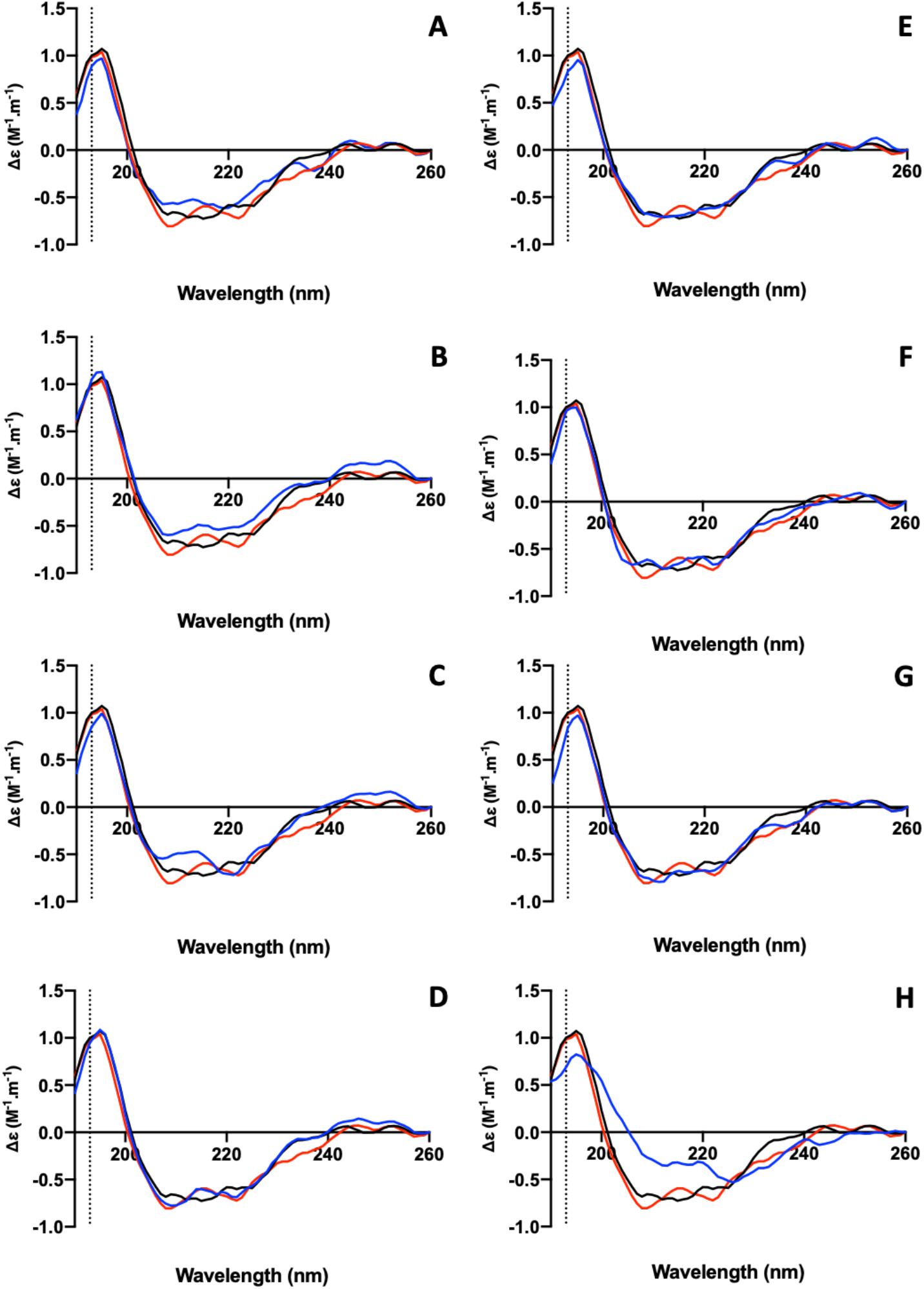
Circular dichroism spectra of nCovS1RBD alone (black) with heparin (red) and chemically modified heparin derivatives (A - H; blue), in 1x PBS, pH 7.4, between 190 and 260 nm. The vertical dotted line indicates 193 nm. The spectra indicate recordings using modified heparins 2-9, with the following predominant disaccharide repeats: (**A**) IdoA2S-GlcNAc6S; (**B**) IdoA2OH-GlcNS6S; (**C**) IdoA2S-GlcNS6OH; (**D**) IdoA2OH-GlcNAc6S; (**E**) IdoA2S-GlcNAc6OH; (**F**) IdoA2OH-GlcNS6OH; (**G**) IdoA2OH-GlcNAc6OH; (**H**) IdoA2,3S-GlcNS3,6S.

